# SCeQTL: an R package for identifying eQTL from single-cell parallel sequencing data

**DOI:** 10.1101/499863

**Authors:** Yue Hu, Xuegong Zhang

## Abstract

**Summary:** With the development of single-cell sequencing technologies, parallel sequencing the transcriptome and genome is becoming available and will bring us the opportunity to uncover association between genotype and phenotype at single-cell level. Due to the special characteristics of single-cell sequencing data, new method is needed to identify eQTL from single-cell data. We developed an R package SCeQTL that uses zero-inflated negative binomial regression to do eQTL analysis on single-cell data. It can distinguish two type of gene-expression differences among different genotype groups. It can also be used for finding gene expression variations associated with other grouping factors like cell lineages or cell types.

**Availability:** The R package is available at https://github.com/XuegongLab/SCeQTL/.

## 1. Introduction

Expression quantitative trait locus or eQTL analysis is an important approach for studying the association and the underlying regulation relationship between variations in the genotype and gene expression. Technologies that can sequence both genome and transcriptome of single cells have been developed recently (Macaulay *et al*., 2015; Dey *et al*., 2015). These technologies give us an opportunity to uncover the association between genetic variations and genes expression at single-cell level, which can help reveal detailed gene regulation mechanisms in processes like tumorigenesis and cell differentiation.

Methods for identifying eQTLs have been well studied for microarray data and bulk RNA-seq data. Typical methods of eQTL mapping include linear regression and ANOVA, where the expression level is taken as the dependent variable and the genotype at a single-nucleotide variation (SNV) site is the explaining factor (Shabalin, 2012; Gatti *et al*., 2009). Among all these methods, most of them are based on the assumption that expression levels or its logarithms follow normal distribution, or Poisson distribution or negative binomial distribution (Sun, 2012). The Krux method used a non-parametric way to identify eQTL and claim their method is more robust (Qi *et al*., 2014). These existing methods including the non-parametric one will lose their power when applied on single-cell RNA-seq data because of the excess of zero values.

The phenomenon of excess of zero values is common in single-cell RNA-seq (scRNA-seq) data (Miao and Zhang, 2016). There are mainly two reasons (Miao et al, 2018). Because the amount of total RNAs in a single cell is extremely small, there is high probability that the reverse transcription step may miss some transcripts, causing the expression of these genes not observed in the sequencing data. This is usually called “drop-out” events. Another reason is that gene expression is a stochastic process at single-cell level (Munsky et al 2012). This results in high variation of gene expression between cells. Studying such heterogeneity is one of the major purposes of single-cell sequencing. Because of these special properties, when we analyze eQTLs on scRNA-seq data, we face two possible types of differences among different genotypes: differences in the proportion of zero-values and differences in non-zero expression levels. We call them as “zero-ratio difference” and “expression level difference”, respectively, for convenience. We developed SCeQTL to analyze these two types of differences that may be associated with genotype variations. The method can also be applied to analyze associations of gene expression with other types of groupings such as cell lineages or cell types.

## 2. Zero-inflated generalized linear model

We model the generation of scRNA-seq data as two processes. One is that transcripts are captured in the sequencing and the corresponding gene gets non-negative expression values. The other is that transcripts are missed or the gene is not expressed in the cell, which will result in zero expression values. The second process causes scRNA-seq data to have excess zero values. We find non-zero parts of scRNA-seq data fit the negative binomial distribution well similar to bulk RNA-seq data, but there can be a high probability of a gene being dropped out in the single-cell data.

Therefore, we use a zero-inflated negative binomial regression to model the scRNA-seq data. For gene expression *G* and genotype *S*, there is probability *p* that transcript is dropped out and probability 1 – *p* that gene expression is generated from a negative binomial distribution. We write these as

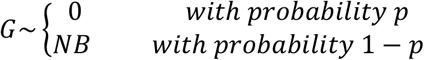

In the equation, μ and θ are the mean and shape parameter of the negative binomial distribution. We call a SNV to be an eQTL of a gene if the probability *p* and/or the mean of negative binomial distribution μ is significantly correlated with the genotype of the SNV in a way that

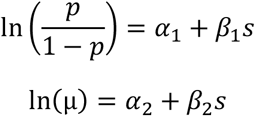

where, parameters *α*_1_, *α*_2_, *β*_1_, *β*_2_ and the shape parameter θ are to be estimated from the data. Using maximum likelihood method to estimate the parameters, we get the log-likelihoods of the full model that includes the genotype as the explaining factor (β_1_ ≠ 0 or β_2_ ≠ 0) and of the reduced model that does not include the genotype (β_1_ = 0 or β_2_ = 0).

According to the generalized linear model theory (Nelder and Baker, 1972), the deviance, which is -2 times the log-likelihood ratio of the reduced model compared to the full model, follows an approximate chi-square distribution with *k* degree of freedom. The *k* is the difference between parameter numbers of the full model and the reduced model. We use the deviance as the test statistic to test for whether *β*_1_ = 0 or *β*_2_ = 0. By these two hypothesis tests, we can identify whether the gene have association with the genotype and what kind of association it is.

## 3. Application examples

Single-cell parallel sequencing data are currently only available in very few labs and are not widely available yet. We used a semi-simulated data to test the power of method proposed. We used the human preimplantation embryos scRNA-seq data (Petropoulos et al., 2016) as the real gene expression data, and split the samples into three groups according to the embryonic day (E5-E7) to mimic three groups of genotypes.

We first checked whether the non-zero data were fitted well with our model using the ‘checkdist’ function in our package. We randomly picked some genes and drew Q-Q plot to compare gene expression distribution with negative binomial distribution. The example of Fig. 1(a) showed that all genes fit the distribution well.

Next, we applied SCeQTL and the widely used MatrixEQTL (Shabalin, 2012) on the data. We found that results of the two methods largely overlapped but there were some cases on which SCeQTL worked better. Fig. 1(b) shows an example that non-zero part had significant difference but MatrixEQTL didn’t find it. One reason is that the negative binomial distribution fit real data better than normal distribution. On the other hand, the zero values in scRNA-seq data caused the means of the three groups to be almost equal, so that MatrixEQTL could not detect the difference. Figure 1(c) gives an example that zero ratio have significant difference but non-zero part is almost same. MatrixEQTL can’t find differences of this type, too.

**Figure 1.**
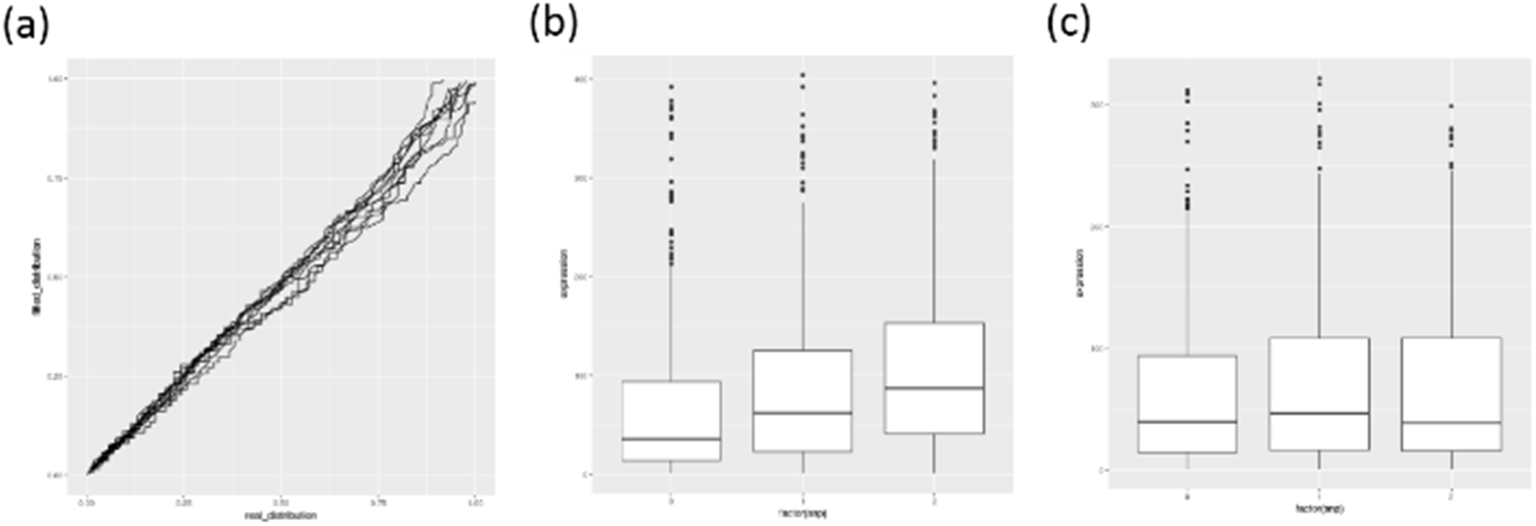
(a) Normalized Q-Q plot for several randomly picked genes (b) An example that non-zero part of gene expression have significant differences among genotype groups, with the two p-values of 9.37 × 10^-9^ and 0.002, respectively. (c) The non-zero part of an example of significant zero-ratio differences. Zero-ratio of three genotype groups are 0.76, 0.26 and 0.26, respectively. The two p-values are 1.09 × 10^-45^ and 0.004, respectively. We can see from the figure that the non-zero expressions are not associated with the genotype, but the drop-out event is.

## 4. Discussion

SCeQTL is a comprehensive software package for eQTL analysis on single-cell genomic and transcriptomic parallel sequencing data. It can also be applied on tasks of finding the association of gene expression with other grouping factors. It provides an effective tool for exploring potential regulatory relationships at single-cell level.

A limitation of the method is that the computation cost is relatively high if applied for eQTL analysis at whole-genome scale. It can take a few minutes on a single computing node to analyze a few hundred gene-SNV pairs. This is mainly due to the iterative procedures in estimating the parameters. However, for most single-cell studies, the cells are from the same tissue sample or closely related samples. We can expect that the number of SNVs among the cells that need to be studied for eQTL analysis is not too large to make the speed of SCeQTL a severe issue in practical applications.

## Supplementary Materials

### Data Preprocessing

Firstly, we remove the effect of the library size. We use the DEseq (Anders and Huber, 2010) way of normalization. That is, the median of the ratios of observed counts is used to measure the sequencing depth.

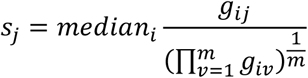

*g_ij_* is the expression level of gene i in sample j. The denominator is calculated by calculating the geometric mean across non-zero samples. As the paper said, this method is more robust than just taking the sum of all genes as sequencing depth since the high expressed gene will dominant the result, which is often seen in single-cell gene expression data. All the samples are normalized by the size factor, and we round down the result to suit our count model.

Next, we remove the genes and variants which are not suitable for the analysis. Genes which are not expressed are removed, genes whose read counts are less than a threshold are treated as not expressed (by default, <=1). We only consider genes whose variances are greater than a threshold (by default, >=5). For variants, only variants creating at least two genotypic groups, each genotype present in at least five samples, are further considered.

When we enter the iteration of analyzing every gene-variant pair, pair that don’t have enough nonzero values (by default, <=5) in one genotype is reported. The estimation of distribution parameter can be far away from true value in this situation. And we find that in real data, there are samples whose expression level is much higher that the others, if we include these samples into consideration, the mean of negative binomial distribution will be overestimated. So we treat these samples as outlier and use robust z score to remove them (by default, >=4). MAD stands for the median absolute deviation.

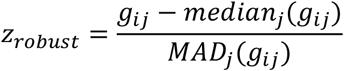

### Parameter estimation

Package ‘pscl’ (Zeileis, Kleiber and Jackman, 2008) is used to estimate the parameter and calculate the log-likelihood now. The package use EM algorithm or BFGS algorithm to iterate updating the parameter.

### Covariates correction

It is common that some hidden covariates may exist in the sampled population, such as age, gender, or other clinical variables. It is important to remove the effect of them from the eQTL study, as otherwise a high percentage of results will be false discoveries. SCeQTL allows user to define a covariate vector x as possible confounding factors to be considered in the analysis. With covariate vector x ∈ R^n^, the models become

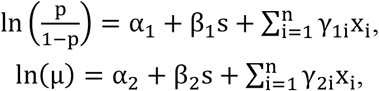

where extra parameter vectors ***γ*** and ***γ_2_*** to be estimated. The hypothesis test process is the same as non-covariates one.

### Multiple test correction

We provide two ways to control the false discovery, Benjamini-Hochberg (BH) method (Benjamini and Hochberg, 1995) and Q-value method. The Q-value method is implement by R package ‘qvalue’ (http://github.com/jdstorey/qvalue). Since several publications come up with other methods for multiple test correction in eQTL mapping (Degnan et al, 2008; Peterson et al, 2016), users can select whether p-value or false discovery rate is returned and the threshold for either of them.

### Experiment result

Figure S1 and S2(a) shows that negative binomial distribution is appropriate for modeling the nonzero part of the data, while the lines in Q-Q plot of other distributions are far away from the diagonal. Figure S2(b) shows that the drop-out event is very common in single-cell RNA-seq data and needs to be considered. Figure S3 show the P value distribution under null hypothesis, we randomly generate the genotype and permutate it and use two methods to calculate the P value. P value distribution of SCeQTL is closed to uniform distribution between 0 and 1, while the result of Matrix eQTL have strong bias.

**Figure S1:**
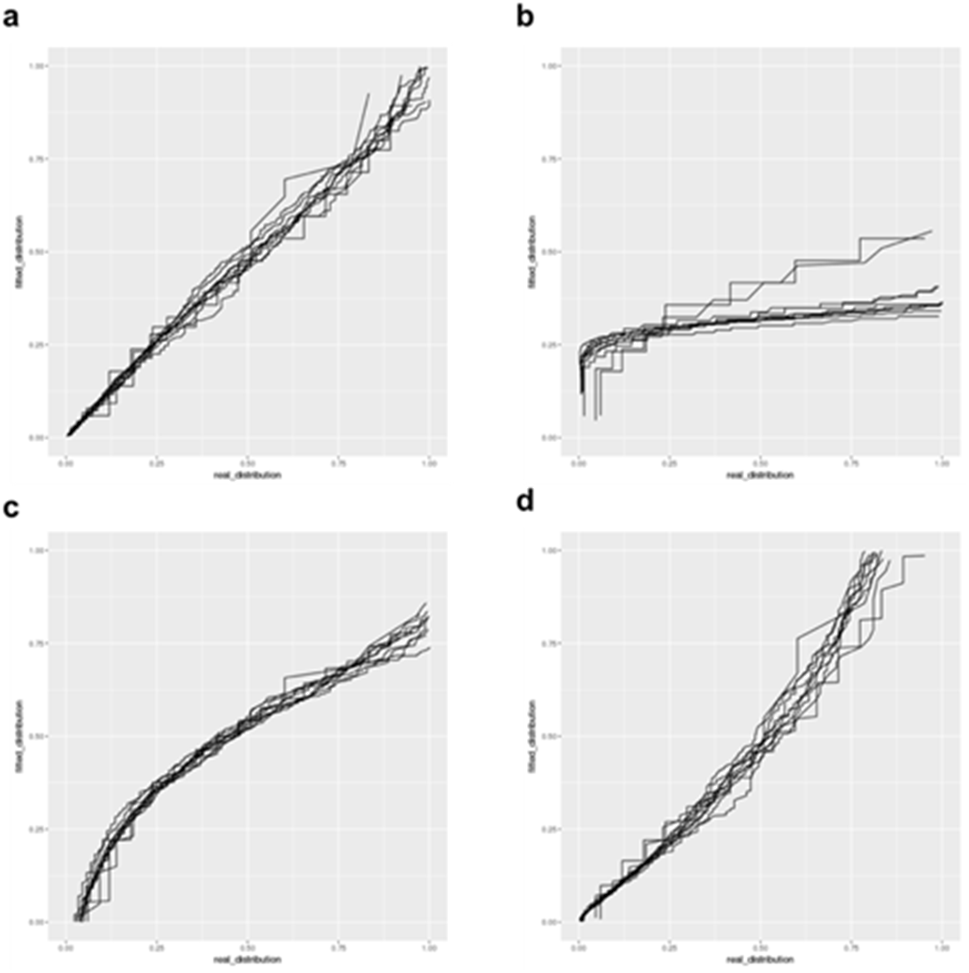
Q-Q plot of (a) negative binomial distribution (b) Poisson distribution (c) Normal distribution (d) log-normal distribution.

**Figure S2:**
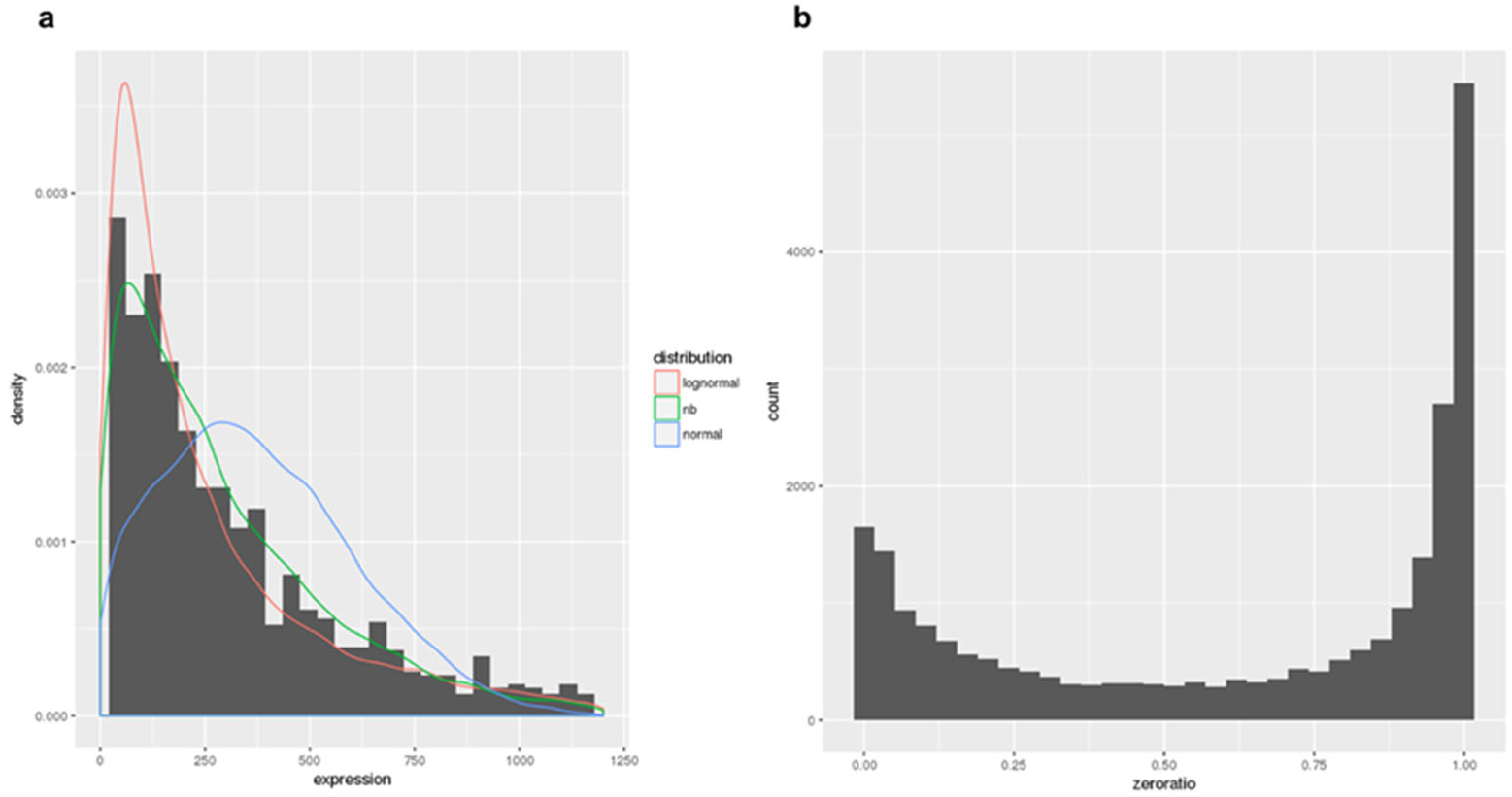
(a) Probability density function of fitted distribution and the histogram of a sample; (b) Histogram of zero ratio of all the genes.

**Figure S3:**
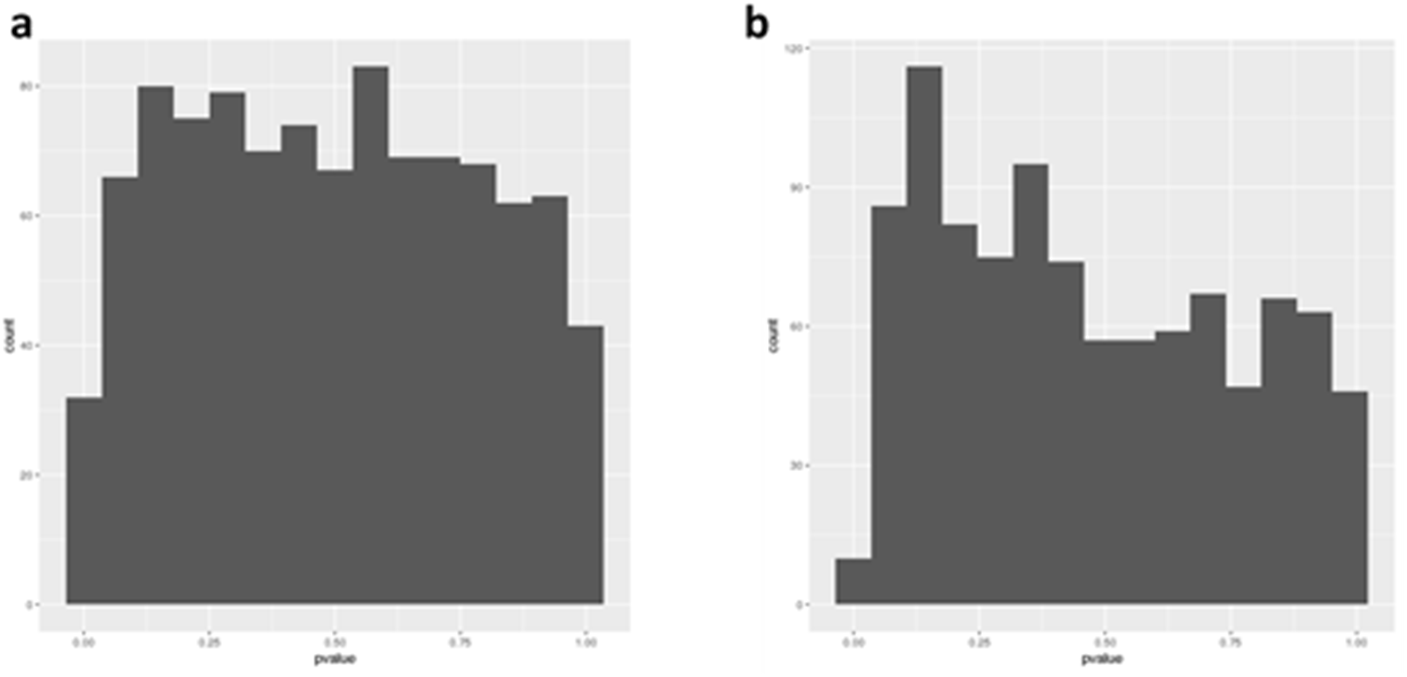
P value distribution under null hypothesis of (a) SCeQTL and (b) Matrix eQTL.

We also found SCeQTL could successfully find some biological interesting results. By dividing the samples by cell lineage, we applied SCeQTL to find genes that vary among different cell lineage. Among all 20,000 genes, SCeQTL found about 20 genes P value less than 10^-40^, 70 genes P value less than 10^-30^ and 200 genes P value less than 10^-20^. In these genes that are significantly correlated with cell lineage, we found some genes have been reported as lineage specific genes. For example, EPI (epiblast) specific genes PRDM14, GDF3, TDGF1, NODAL, SOX2, NANOG, P value of these genes are 5.7 × 10^-45^, 4.7 × 10^-20^, 6.5 × 10^-16^, 4.0 × 10^-21^, 5.1 × 10^-37^, 4.8 × 10^-25^; TE (trophectoderm) specific genes GATA2, GATA3, DAB2; P value of these genes are 4.1 × 10^-20^, 6.7 × 10^-26^, 3.3 × 10^-21^; PE (primitive endoderm) specific genes HNF1B, PDGFRA, GATA4, P value of these genes are 5.7 × 10^-32^, 1.7 × 10^-23^, 2.0 × 10^-35^. All these lineage specific genes rank top 200 in our result. What’s interesting is that quite a lot of these genes have obvious zero ratio differences, which may prove that zero ratio differences are important as non-zero part differences in single cell data since genes in single cell may close due to some reasons.

We also manually checked several genes that have not been reported. We found most significant genes truly have difference among groups. Figure S4 shows three examples. The results need further investigations.

**Figure S4:**
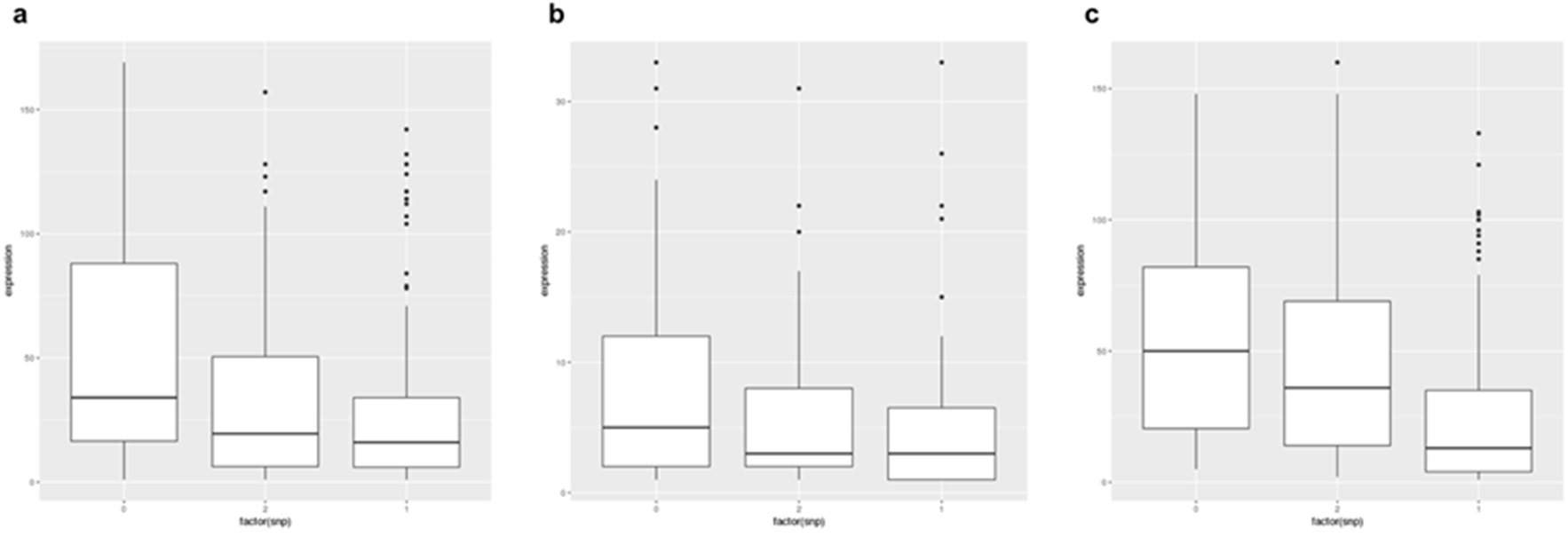
Figure S4: Examples of lineage differential expressed genes (a) PSORS1C2, Zero-ratio of three genotype groups are 0.18, 0.85, and 0.57, respectively. The two p-values are 4.1 × 10^-50^ (b) PTPRZ1, Zero-ratio of three genotype groups are 0.45, 0.93 and 0.73, respectively. The two p-values are 4.8 × 10^-33^ (c) AMDHD1, Zero-ratio of three genotype groups are 0.60, 0.79, and 0.38, respectively. The two p-values are 1.5 × 10^-23^.

